# RNA-DNA triplex-forming miRNAs define an evolutionarily recent chromatin regulatory mechanism

**DOI:** 10.64898/2026.02.25.707792

**Authors:** Michell Martin, Consuelo Jalife, Catalina Méndez, Cristian Montecinos, David A. Brown-Brown, Francisca Díaz-Tejeda, Valeria Caffi, Kattina Zavala, Fabian Kugler, Loreto Molina-González, Guillermo Valenzuela-Nieto, Gernot Längst, Gonzalo A. Mardones, Roberto Munita, Juan C. Opazo, Rodrigo Maldonado

## Abstract

MicroRNAs (miRNAs) are best known for their role in post-transcriptional gene regulation in the cytoplasm. However, a subset of miRNAs has been detected in the nucleus, suggesting additional regulatory functions. Here, we systematically characterize chromatin-associated small non-coding RNAs in the human pancreatic cancer cell line PANC-1. Using chromatin RNA immunoprecipitation coupled with small RNA sequencing, we show that the chromatin-associated small RNA population differs markedly from the bulk nuclear RNA pool and is strongly enriched in miRNAs. Among these, miR-21 represents the most abundant chromatin-associated species. Sequence analyses revealed that a subset of these miRNAs fulfills the requirements for RNA-DNA triplex formation at genomic regulatory regions. Gel-shift assays further demonstrate that Argonaute2 (Ago2) directly interacts with triple-helical nucleic acid structures *in vitro*, suggesting a potential mechanistic link between triplexes and Ago2-chromatin engagement. Evolutionary analyses indicate that these triplex-forming chromatin-associated miRNAs are largely restricted to anthropoid primates, in contrast to broadly conserved non-triplex-forming miRNAs. Together, our results identify a population of chromatin-associated miRNAs and provide evidence for a potential structural mechanism linking miRNAs, Ago2, and chromatin.

## Introduction

Chromatin regulation involves a complex interplay between DNA, histones, and regulatory proteins that together shape gene expression programs. In addition to these canonical components, increasing evidence indicates that non-coding RNAs (ncRNAs) contribute to chromatin organization and transcriptional control. A substantial fraction of cellular RNAs remains associated with the nucleus and chromatin rather than being exported to the cytoplasm, suggesting that RNA molecules are integral elements of the chromatin regulatory environment. Indeed, chromatin-bound RNAs have been implicated in diverse processes including chromatin accessibility, heterochromatin maintenance, and genome stability (1–4).

Among the best-characterized histone modifications that mark transcriptionally active chromatin are H3K27 acetylation (H3K27ac) and H3K4 trimethylation (H3K4me3). H3K27ac is a histone mark associated with active enhancers and active gene regulatory regions, deposited by acetyltransferases such as CBP/p300 in response to transcriptional activation signals (5). H3K4me3 is enriched at the transcription start sites of active genes, where it facilitates the recruitment of the basal transcription machinery and chromatin remodeling complexes (6). Both marks therefore serve as reliable indicators of transcriptionally permissive chromatin environments. Because RNA-DNA triplex formation has been proposed to be favored in open, nucleosome-proximal chromatin contexts (7), these active marks provide an appropriate framework for enriching chromatin fractions where triplex-mediated regulatory interactions are most likely to occur.

MicroRNAs (miRNAs) are a class of small non-coding RNAs (sncRNAs) best known for their role in post-transcriptional gene regulation in the cytoplasm, where they associate with Argonaute proteins to form the RNA-induced silencing complex (RISC) (8). However, accumulating evidence indicates that a subset of miRNAs localizes to the nucleus and can influence transcriptional processes. Nuclear miRNAs have been shown to modulate gene expression by targeting promoter or enhancer regions in an Argonaute-dependent manner, leading either to transcriptional repression or activation (9–11). These observations suggest that miRNAs may participate directly in chromatin-based regulatory mechanisms.

One proposed mechanism by which miRNAs could interact with genomic DNA involves the formation of RNA-DNA triplex structures (12). Triplexes arise when a third nucleic acid strand binds to the major groove of double-stranded DNA through Hoogsteen or reverse Hoogsteen hydrogen bonding (13,14). These interactions are highly sequence dependent and have been described for several ncRNAs that regulate gene expression by targeting specific genomic loci (15–18). Computational and experimental studies have suggested that a subset of miRNAs may also possess sequence features compatible with triplex formation, potentially allowing them to directly associate with chromatin regulatory elements (7,12,19). Triplex-forming sequences are identified based on the ability of a single-stranded RNA (ssRNA) or DNA (ssDNA) oligonucleotide (the triplex-forming oligonucleotide, TFO) to bind, via Hoogsteen or reverse Hoogsteen hydrogen bonds, to the purine-rich strand of a polypurine–polypyrimidine double-stranded DNA (dsDNA) sequence (the triplex target site, TTS). This interaction requires the TFO to satisfy specific nucleotide composition constraints, primarily purine-rich or pyrimidine-rich character, and a minimum length of approximately 13–15 nucleotides of continuous complementarity (13–15). Despite these observations, the identity and functional properties of chromatin-associated miRNAs remain poorly defined. In particular, it is unclear whether miRNAs that associate with chromatin possess distinct sequence characteristics, how they might interact with chromatin-associated protein complexes, and whether these mechanisms have emerged during evolution as specialized regulatory strategies.

Here, we systematically characterize chromatin-associated small RNAs in the human pancreatic cancer cell line PANC-1. Using chromatin RNA immunoprecipitation followed by small-RNA sequencing, we identify a chromatin-associated RNA population that is distinct from the bulk nuclear RNA pool and is strongly enriched in miRNAs. Among these, miR-21 emerges as the most abundant chromatin-associated species. We further identify a subset of miRNAs with predicted RNA-DNA triplex-forming potential and demonstrate that Ago2 interacts with triple-helical nucleic acid structures *in vitro*. Finally, evolutionary analyses reveal that triplex-forming chromatin-associated miRNAs are largely restricted to anthropoid primates. Together, our results suggest that triplex-mediated miRNA-chromatin interactions represent an evolutionarily recent mechanism linking miRNAs and Argonaute proteins to chromatin regulation.

## Material and Methods

### Cell culture

PANC-1 and HeLa cells were cultured in Dubecco’s modified Eagle Medium (DMEM) supplemented with glucose (4.5 g/L), L-glutamine (584 mg/L), sodium pyruvate (110 mg/L), and 10% fetal bovine serum. Cells were maintained at 37 °C in 5% CO_2_ and subcultured using 0.25% trypsin.

### Nuclear preparations

PANC-1 cells grown in 150-mm culture dishes to ∼80% confluency were washed and incubated with 10 mL of 1X PBS containing 20 µL of 500 µM Thiazole Orange (390062 Sigma-Aldrich, prepared in DMSO) for 5 min at room temperature with agitation. Cells were crosslinked with 1% formaldehyde in 1X PBS for 8 min at room temperature, and the reaction was quenched with 125 mM glycine for 5 min. Cells were then scraped, transferred to 1.5 mL microcentrifuge tubes, and washed with ice-cold 1X PBS supplemented with RNase and protease inhibitors. For subcellular fractionation, cell pellets were resuspended in 850 µL of 1X PBS containing 0.2% Triton X-100 and supplemented with RNase and protease inhibitors. Cells were lysed by 20 pipetting cycles using a P1000 pipette, followed by vortexing for 30 sec. An aliquot of 50 µL was collected as the Whole Cell Extract (WCE). Samples were then centrifuged at 12,000 × g for 30 sec at 4°C. The supernatant was collected as the cytoplasmic fraction, while the pellet corresponded to the nuclear fraction. Nuclear pellets were resuspended in 500 µL of 1X PBS containing 0.2% Triton X-100 and RNase and protease inhibitors, divided equally into two aliquots of 250 µL, and centrifuged again at 12,000 × g for 30 sec at 4 °C. Supernatants were discarded, and nuclear pellets were stored at -80 °C until further processing.

### Enzymatic digestion of chromatin

Frozen nuclear pellets were thawed and resuspended in digestion buffer (15 mM Tris-HCl, 60 mM KCl, 15 mM NaCl, 5 mM MnCl2, 0.5 mM EGTA, 300 mM Sucrose, 0.5 mM 2-mercaptoethanol, pH 7.4), followed by DNase I digestion for 5 min at 37 °C. The reaction was stopped by 50 mM EDTA addition, and samples were adjusted to 600 µL with RIPA buffer (10 mM Tris-HCl, 1 mM EDTA, 0.1% SDS, 0.1% sodium deoxycholate, 1% Triton X-100, pH 7.6) and homogenized at 4 °C for 15 min before ChRIP assays. For DNA fragment analysis, crosslinks were reversed by overnight incubation at 65 °C, followed by RNase A treatment (1 h at 37 °C) and proteinase K digestion (45 min at 50 °C). DNA was precipitated by adding ammonium acetate (0.5 volumes, 7.5 M, pH 7.7), ethanol (2 volumes), and glycogen (20 µg), incubating at -20°C, centrifuging, washing with 75% ethanol, and resuspending in nuclease-free water. DNA concentration was measured using the Qubit dsDNA HS assay. Fragment size distribution was assessed by electrophoresis on 1.3% agarose gels in TBE buffer (89 mM Tris pH 7.6, 89 mM Boric Acid, 2 mM EDTA), revealing DNA fragments predominantly in the 200–500 bp range.

### Western Blot

Proteins resolved by SDS-PAGE were transferred to methanol-activated PVDF membranes at 100 V for 1 h at 4 °C. Transfer efficiency was verified by Ponceau S staining. To assess cellular fractionation, membranes were probed for α-tubulin (T5168, Sigma-Aldrich) and histone H3 (ab1791, Abcam) as cytoplasmic and nuclear markers, respectively. For chromatin immunoprecipitation assays, membranes were incubated with anti-H3, anti-H3K27Ac (ab4729, Abcam), and anti-H3K4me3 (ab8580, Abcam). Membranes were blocked in TBS 0.1% Tween 20 (TBS-T) containing 5% non-fat milk for 1 h at room temperature, then incubated overnight at 4 °C with the corresponding primary antibodies. Afterwards, membranes were washed 5 times with TBS-T and incubated with HRP-Protein A secondary reagent for 1 hr at room temperature. Signals were detected using chemiluminescence on a Syngene G:Box imaging system.

### Chromatin-RNA Immunoprecipitation

Chromatin-RNA immunoprecipitation was performed using enzymatically digested PANC-1 nuclear extracts. Briefly, 600 µL of digested chromatin were pre-cleared with 20 µL of Dynabeads Protein G magnetic beads (Invitrogen 10004D) equilibrated in RIPA buffer for 1 h at 4 °C with rotation to reduce nonspecific binding. After magnetic separation, the supernatant was recovered as the pre-cleared sample. Aliquots corresponding to 5% of the material were reserved as input controls for RNA purification and Western blot analysis and stored at -80 °C. For immunoprecipitation, pre-cleared chromatin was incubated overnight at 4 °C with 4 µg of the corresponding antibodies under constant rotation. Magnetic beads (20 µL, pre-washed in RIPA buffer) were then added and incubated for 1 h at room temperature with rotation. Beads were collected magnetically, and a fraction of the unbound supernatant was retained for Western blot analysis. Beads were sequentially washed five times (two washes with RIPA, one wash with high-salt RIPA containing 0.5 M NaCl, and two additional washes with RIPA). Immunoprecipitated material was resuspended in a proteinase K buffer (10 mM Tris-HCl, 10% SDS, 50 mM EDTA, pH 7,4). Five percent was reserved for Western blot validation, and the remaining material was processed for RNA extraction. RNAs were extracted from immunoprecipitated and input samples using TRIzol™ reagent according to the manufacturer’s instructions, with glycogen as a carrier. Total RNA was precipitated, washed with 75% ethanol, and resuspended in nuclease-free water. Enrichment of small RNAs was performed using the mirVana™ miRNA Isolation Kit, and RNA concentrations were determined using the Qubit™ RNA HS Assay Kit.

### Small RNA sequencing and analysis

Small RNAs were first subjected to quality control to assess RNA integrity and small RNA concentration using the Qubit™ RNA HS Assay Kit and capillary electrophoresis (Figure 1D). Samples meeting the required quality and concentration criteria were processed for small RNA library preparation using the QIAseq™ miRNA Library Kit (Qiagen), following the manufacturer’s instructions. Library quality and size distribution were evaluated before sequencing. Prepared libraries were sequenced at Austral-*omics* (Valdivia, Chile) using a NextSeq 550 platform (Illumina).

**Figure 1.**
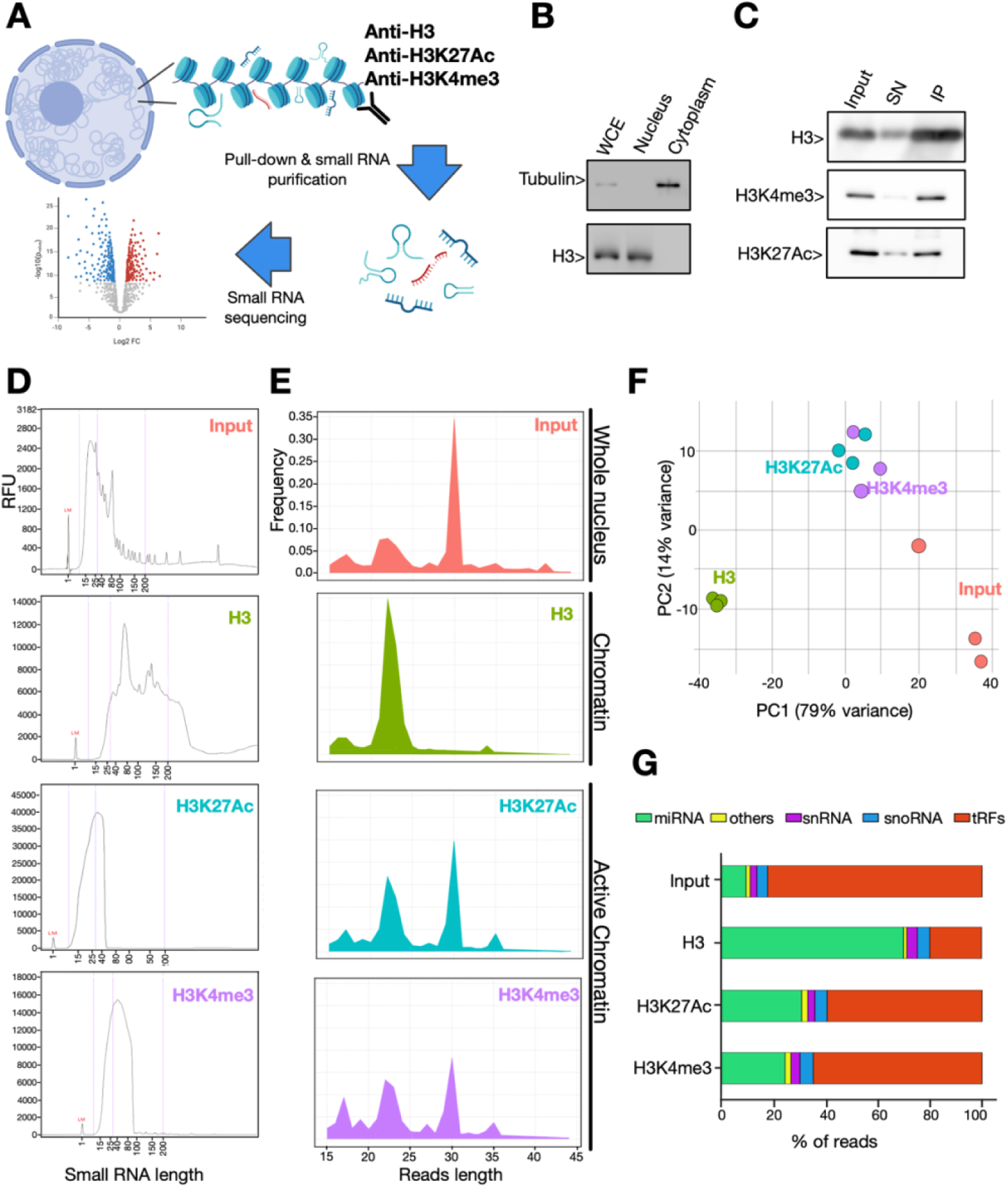
Differential distribution of small RNAs in the nucleus and chromatin in PANC-1 cells. A) General overview of the chromatin-RNA immunoprecipitation (ChRIP) workflow from PANC-1 nuclear extracts. For a detailed version check the Supplementary Figure 1A. B) Subcellular fractionation of PANC-1 cells assessed by Western blot (WB) analysis of whole-cell extract (WCE), nuclear, and cytoplasmic fractions. α-Tubulin and histone H3 were used as cytoplasmic and nuclear markers, respectively. C) WB validation of H3, H3K27ac, and H3K4me3 immunoprecipitations showing input, non-bound supernatant (SN), and immunoprecipitated (IP) fractions. D) Size distribution profiles of small RNAs purified from each immunoprecipitated chromatin fraction. E) Size distribution of the sequencing reads after analysis. F) Principal component analysis (PCA) of small RNA-sequencing datasets. G) Percentage distribution of RNA biotypes among differentially enriched small RNAs.

Raw read quality was assessed using FastQC (20) and MultiQC (21). UMIs (12 nucleotides) were extracted using UMI-tools (22) with the pattern .+(?P<DISCARD_1>AACTGTAGGCACCATCAAT){s<=2}(?P<UMI_1>.{12}), followed by adapter removal using Cutadapt (23). Reads shorter than 15 nucleotides or containing ambiguous bases were discarded. Reads were aligned against an rRNA reference using Bowtie2 (24). Non-rRNA reads were aligned against a curated human small non-coding RNA reference derived from HGNC and obtained from RNAcentral (release 20.0; hgnc.fasta) using Bowtie2. PCR duplicates were removed using UMI-tools (dedup, --method=directional). Reads were classified into RNA biotypes (tRNA, miRNA, snRNA, snoRNA, rRNA) using a custom Python script with HGNC annotations. Relative biotype abundance and read length distributions were calculated for each condition (H3, H3K27Ac, H3K4me3, Input). Differential enrichment analysis was performed using DESeq2 (25), filtering features with <10 total reads. Pairwise comparisons between histone marks and Input identified differentially enriched transcripts (FDR < 0.05, |log2FC| > 1). Visualization was performed using ggplot2 and EnhancedVolcano (26).

### Real time PCR

Total RNA (1 µg) from PANC-1 or HeLa cells was reverse-transcribed using the RevertAid RT Kit (Thermo Scientific) following the manufacturer’s instructions for mRNA analysis. For miRNA quantification, miR-21-5p expression was measured using a stem-loop reverse transcription (5’-CACCGTTCCCCGCCGTCGGTGTCAACA-3’) and quantitative PCR approach (Forward 5’-CCCGCCTAGCTTATCAGACTG-3’, Reverse 5’-GCCGTCGGTGTCAACATCA-3’), as previously described (27), which enables specific detection of mature miRNAs. Specific primers against β-Actin (*ACTB*) mRNA (Forward 5’-ACCACCATGTACCCTGGCATT-3’, Reverse 5’- CCACACGGAGTACTTGCGCTCA-3’) were used as an internal normalization control (27). Quantitative PCR reactions were carried out using SsoAdvanced™ Universal SYBR® Green Supermix (Bio-Rad, Cat. No. 1725271) and analyzed on a Rotor-Gene RG3000 thermal cycler (QIAGEN).

### Cloning, expression, and purification of recombinant proteins

Purified recombinant wild-type Ago2 was kindly provided by Prof. Dr. Gunter Meister (Regensburg Center for Biochemistry, Laboratory of RNA Biology, University of Regensburg). Recombinant constructs encoding the N-Terminal (MYSGAGPALAPPAPPPPIQGYAFKPPPRPDFGTSGRTIKLQANFFEMDIPKIDIYHYELDIKPEKCPRRVNREIVEHMVQHFKTQIFGDRKPVFDGRKNLYTAMPLPIGRDKVELEVTLPGEGKDRIFKVSIKWVSCV) and PIWI (PQGRPPVFQQPVIFLGADVTHPPAGDGKKPSIAAVVGSMDAHPNRYCATVRVQQHRQEIIQDLAAMVRELLIQFYKSTRFKPTRIIFYRDGVSEGQFQQVLHHELLAIREACIKLEKDYQPGITFIVVQKRHHTRLFCTDKNERVGKSGNIPAGTTVDTKITHPTEFDFYLCSHAGIQGTSRPSHYHVLWDDNRFSSDELQILTYQLCHTYVRCTRSVSIPAPAYYAHLVAFRARYHLVDKEHDSAEGSHTSGQSNGRDHQALAKAVQVHQDTLRTMYFA) domains of human Ago2 were cloned into the pGEX-TEV expression vector, generating GST-fusion proteins. Plasmids were transformed into *Escherichia coli* DH5α competent cells by heat shock, and transformants were selected on Luria-Bertani agar plates supplemented with ampicillin (100 µg/mL). Positive colonies were screened by colony PCR using domain-specific primers, and plasmid DNA was purified using the E.Z.N.A.® Plasmid DNA Kit (Omega Bio-Tek). Insert integrity was confirmed by Sanger sequencing.

For protein expression, verified plasmids were transformed into *E. coli* B834(DE3)pLysS cells. Small-scale induction assays were performed to optimize expression conditions. Briefly, cultures were grown in Luria-Bertani medium containing ampicillin (100 µg/mL) at 37 °C to mid-log phase and induced with IPTG (1 mM for GST-N-terminal, and 0.33 mM for GST-PIWI). Protein expression was carried out at 25 °C for 16 h with shaking. Bacterial cells were harvested by centrifugation and resuspended in lysis buffer (50 mM Tris-HCl, 500 mM NaCl, 5 mM EDTA, 1 mM DTT, pH 8.0) supplemented with 2 mM PMSF. Cells were lysed by sonication, and the insoluble material was removed by centrifugation. Clarified lysates were subjected to chromatography on Glutathione Sepharose 4B (Cytiva Cat. nr. 170754), equilibrated in lysis buffer, and washed extensively with the same buffer. Elution of GST-fusion proteins was performed with elution buffer (50 mM Tris-HCl, 150 mM NaCl, 5 mM EDTA, 1 mM DTT, 20 mM reduced glutathione, pH 8.0). Eluted fractions were analyzed by SDS–PAGE to assess purity and yield. Large-scale purifications were scaled-up using 3-4 L of bacterial culture under the same optimized conditions. Affinity-purified GST-fusion proteins were further purified by size-exclusion chromatography using a HiLoad 16/600 Superdex 200 pg column (Cytiva Cat. nr. 90100040) equilibrated in TA buffer (40 mM Tris, 10 mM CH_3_COONa, 10 mM MgCl_2_, adjusted to pH 7,4 with glacial acetic acid). Chromatography was performed on an ÄKTA Prime Plus system at 4 °C, with absorbance monitored at 280 nm. Fractions (2.5 mL) corresponding to major elution peaks were collected and analyzed by SDS–PAGE to identify fractions containing the recombinant Ago2 domains. Peak fractions were pooled and stored at -80 °C for further assays.

### Triplex predictions

TTS were identified using PATO (28) with lengths ranging from 13 to 29 bp, no mismatches in the polypurine motif, and were filtered to ensure a minimum distance of 200 bp between adjacent sites. For each TTS, all potential Hoogsteen base-pairing motifs were generated to construct the TFO database used for the BLAST searches.

Triplex-forming oligonucleotides (TFOs) corresponding to expressed small RNAs were identified using BLAST (29) searches against a database of all possible TFOs that perfectly match defined human triplex target sites (TTS) via Hoogsteen base pairing. BLAST searches were performed with blastn in blastn-short (-task “blastn-short”) mode, restricted to the plus strand, requiring 100% sequence identity (-perc_identity 100). BLAST hits were subsequently filtered to retain only alignments with a minimum alignment length of 13 nucleotides.

### Electrophoretic Mobility Shift Assays

Triplexes were assembled as previously described (17). Briefly, 50 nM Cy5-labeled triplex target sites (TTS, double-stranded DNA forward 5’ Cy5-TCT TTT TTT TTT TTT TTC TTT TTT CCT CCT TTT TTT TTC C-3’, and reverse 5’-GGA AAA AAA AAG GAG GAA AAA AGA AAA AAA AAA AAA AAG A-3’) were incubated with 150 nM of the Cy3-labeled triplex-forming oligonucleotides (TFO, single-stranded DNA or RNA 5’ Cy3-ССU UUU UUU UUC CUC CUU UUU UCU UUU UUU UUU UUU UUC U -3’) in TA buffer for 15 min at 37 °C. The self-triplex oligonucleotides (STO) contain the same sequences to form the triplex, but in the DNA form, with a 4 T bases as a linker between the forward/reverse strands and the reverse strand/TFO. The STO were folded under identical conditions to ensure triplex formation. For protein interaction assays, pre-formed triplexes or STO were incubated with increasing concentrations of the indicated proteins for 10 min at 30 °C. Complexes were resolved on native 12% polyacrylamide gels prepared in TA buffer and electrophoresed at 15 V/cm. Gels were imaged by fluorescence detection using a GenoSens 200 Series gel documentation system (CLINX).

### Analysis of the evolutionary distribution and conservation

We annotated the sequences of 14 miRNA genes in representative species of all major groups of jawed vertebrates. Our sampling included mammals, birds, reptiles, amphibians, lungfish, coelacanths, bony fishes, and cartilaginous fishes. Human miRNA sequences were retrieved from RNAcentral (30) and used to identify either the syntenic region (when the miRNA was located in an intergenic region) or the host gene (when the miRNA was embedded within a protein-coding gene) in Ensembl v115 (31). Based on this information, we searched the NCBI database (32) for the corresponding region using tblastn (33), with the amino acid sequence of one of the syntenic genes or the host gene containing the miRNA in humans as the query. Once the target genomic region was obtained in the different jawed vertebrate species, we performed a blast2seq (33) search using the human sequence as the query. After identifying the region of interest in each sampled species, we retrieved the 5p and 3p mature miRNA sequences from miRBase (https://www.mirbase.org/) (34) (Supplementary Table 3). Finally, precursor miRNA nucleotide sequences were aligned using MAFFT v7.49 (35) in Geneious Prime.

## Results

### Chromatin-associated small RNAs are enriched in miRNAs

To investigate the composition of chromatin-associated small RNAs, we performed chromatin-RNA immunoprecipitation (ChRIP) from nuclear extracts of PANC-1 cells followed by small RNA sequencing (Fig. 1A, Supplementary Figure 1A). Cell fractionation efficiency was confirmed by Western blot using tubulin and histone H3 as cytoplasmic and nuclear markers, respectively (Figure 1B). Then, nuclear fraction was treated with DNAseI to digest chromatin, which generated predominantly mono- and di-nucleosomal fragments, indicating suitable conditions for ChRIP analysis (Supplementary Figure 1B). Nuclear extracts were used as input controls, while chromatin fractions were isolated using an anti-histone H3 antibody (Figure 1C, upper panel). Because RNA-DNA triplex formation has been proposed to occur preferentially at transcriptionally active chromatin regions (7), additional ChRIP experiments were performed using antibodies against the active histone marks H3K27ac and H3K4me3 (Figure 1C, middle and lower panels). These four fractions were further subjected to small RNAs purification and profiling, and their size distribution revealed distinct patterns. Nuclear samples were enriched in small RNAs ranging from 15 to 100 nt, whereas whole chromatin samples displayed a broader distribution spanning between approximately 20 and 200 nt (Figure 1D). In contrast, active chromatin fractions exhibited narrower size profiles, with H3K27Ac-associated small RNAs predominantly between 15 and 40 nt, and H3K4me3-associated small RNAs between 15 and 90 nt (Figure 1D). These results indicate that our optimized conditions effectively preserved small RNA-chromatin associations, enabling high-quality samples with specific size distributions suitable for library preparation and sequencing.

Analysis of sequenced reads lengths showed that nuclear extracts were enriched in ∼30 nt RNAs, whole chromatin fractions in 21-24 nt RNAs, and active chromatin fractions showed a bimodal distribution reflecting both populations, revealing a clear difference between nuclear and chromatin-associated small RNA populations (Figure 1E). Principal component analysis (PCA) confirmed high reproducibility across biological replicates and highlighted differences in the profiles between nuclear, whole chromatin, and active chromatin samples. Additionally, PCA showed that the two active chromatin marks yielded highly similar small RNA populations (Figure 1F). Biotype enrichment analysis revealed a predominance of tRNA-derived fragments (tRFs) in nuclear extracts, miRNAs in the whole chromatin fraction, and a mixed population of both in the active chromatin samples (Figure 1G). These results indicate that chromatin-associated small RNAs represent a distinct subset of the nuclear RNA population that is strongly enriched in miRNAs.

### MiRNAs are the most enriched small RNAs, while miR21 is the most abundant in chromatin

ChRIP-seq analysis revealed a differential enrichment of small RNA classes in chromatin compared to the nuclear compartment of PANC-1 cells. To identify the species preferentially associated with chromatin, we compared whole chromatin and active chromatin fractions versus nuclear samples. This analysis showed that a subset of miRNAs was strongly enriched in both, whole and active chromatin, whereas tRNA-derived fragments (tRFs) were preferentially enriched in nuclear extracts relative to chromatin (Figure 2A, Supplementary Table 1). It is important to note that volcano plots (Figure 2A) display differential enrichment ratios between fractions, not absolute expression levels. As small RNAs are generally present at very low absolute levels (compared with coding transcripts) can appear as highly enriched simply by virtue of the mathematical comparison yet remain below biologically meaningful detection thresholds. This distinction is critical for interpreting our data, the high enrichment ratios do not necessarily imply biologically relevant abundance. To address this, we complemented the enrichment analysis with a direct quantification of absolute read counts normalized as counts per million (CPM), which identified miR-21 as the most abundant small-RNA across chromatin-associated fractions, with a marked preference for whole chromatin (Figure 2B, upper plot; Supplementary Table 1). The strong chromatin association of miR-21 is therefore supported both by its relative enrichment and by its high absolute abundance, distinguishing it from other enriched species that are detectable only at minimal read counts. This phenomena remained evident after normalization to pancreatic tissue miRNA expression profiles (36) and nuclear samples (Figure 2B, lower plot).

**Figure 2.**
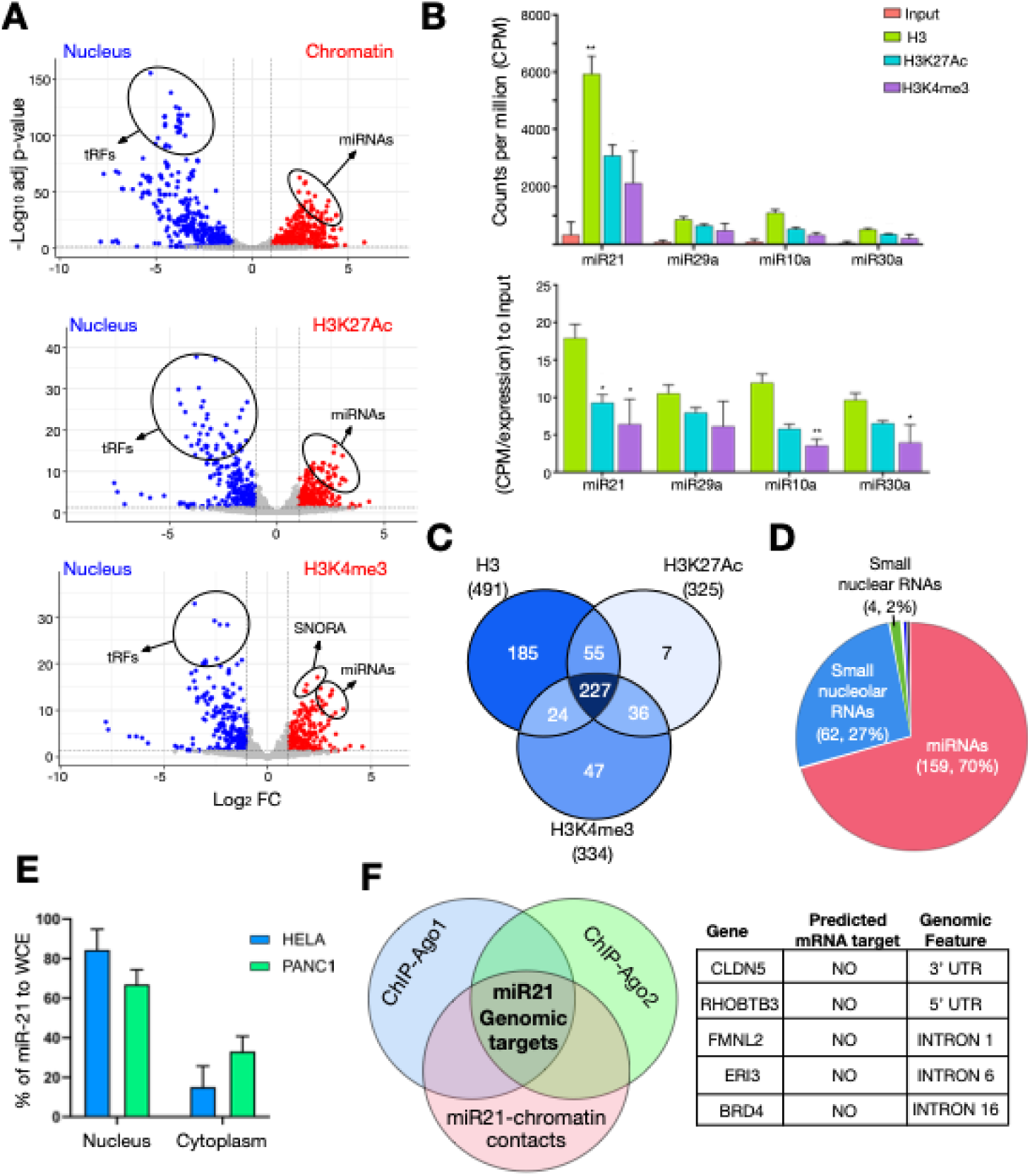
miRNAs are the main small RNAs that associate with chromatin in PANC-1 cells. A) Differential enrichment analysis of small RNA-seq datasets comparing chromatin (H3) and active chromatin (H3K27ac and H3K4me3) fractions against nuclear small RNAs. Volcano plots depict statistical significance (-Log10 Adj p-values) versus fold change (Log2 FC). B) Comparison of miRNA abundance expressed as counts per million (CPM) and CPM normalized to tissue expression relative to the input for the most enriched miRNAs across nuclear and chromatin-associated fractions. Statistical significance is indicated (\**p* < 0.05; **p < 0.01; two-tailed Student’s *t*-test). C) Venn diagram illustrating shared and unique significantly enriched small RNAs in chromatin and active chromatin relative to nuclear fractions. D) Biotype distribution of the 227 small RNAs commonly enriched in chromatin and active chromatin. E) Quantitative real-time PCR analysis showing the relative abundance of miR-21 in nuclear and cytoplasmic fractions normalized to whole-cell extracts from PANC-1 and HeLa cells. F) Left, Venn diagram illustrating putative miR-21 genomic interaction sites inferred from the overlap of Ago1-, Ago2-, and miR-21-associated chromatin regions. Right, schematic representation of potential miR-21 target genes.

Read alignment further revealed that miR-21-5p was the predominant mature miRNA species associated with chromatin (Supplementary Figure 2); therefore, miR-21-5p is referred to as miR-21 throughout this work, unless otherwise specified. Consistent with these findings, comparative analyses showed that approximately 50–60% of chromatin-associated small RNAs were shared between whole and active chromatin fractions (Figure 2C), with miRNAs representing the predominant biotype among these shared species (Figure 2D).

As miR-21 appeared as the most abundant chromatin-associated miRNA, we assessed its nuclear enrichment, compared to the cytoplasm, by quantitative real-time PCR following the same fractionation procedure used for ChRIP, omitting formaldehyde crosslinking. This analysis corroborated our previous results, revealing that miR-21 levels were approximately threefold higher in the nuclear fraction (most probably chromatin) than in the cytoplasmic fraction in both PANC-1 and HeLa cells (Figure 2E).

To descriptively explore a potential association between miR-21 and chromatin-linked transcriptional regulation, we next identified candidate genomic loci occupied by miR-21. This was achieved by integrating published Ago1 and Ago2 chromatin immunoprecipitation datasets (37) with miR-21 chromatin interaction sites retrieved from the RNA-Chrom database (38). This approach identified 90 genomic loci co-associated with Ago1, Ago2, and miR-21 (Supplementary Table 2). Notably, none of the corresponding genes have been reported as miR-21 targets at the mRNA level based on miRBase predicted targets. By further restricting this set to loci containing fully or partially complementary sequences to the miR-21 seed region, we identified five potential candidates as genomic targets (Figure 2F).

Together, these results indicate that the intranuclear distribution of small RNAs in PANC-1 cells is characterized by a selective enrichment of miRNAs on chromatin and provide a descriptive framework for miR-21 association with chromatin for future functional investigations.

### A subset of chromatin miRNAs displays triplex-forming potential

RNA-DNA triplex formation represents a potential mechanism through which RNAs can directly associate with genomic DNA. We therefore evaluated whether chromatin-associated miRNAs possess sequence features compatible with triplex formation.

To identify miRNAs with triplex-forming potential, we used PATO to predict polypurine-polypyrimidine triplex target sites (TTSs) across the human genome, applying a minimum length of 13 bp, no mismatches in the polypurine motif, and a minimum spacing of 200 bp between adjacent sites. For each predicted TTS, all possible TFO sequences compatible with Hoogsteen base pairing were generated and compiled into a searchable database. Chromatin-associated miRNA sequences were then queried against this TFO database using BLAST in short-sequence mode, requiring 100% identity and a minimum alignment length of 13 nucleotides. Comparative analyses using this workflow revealed that approximately 3–6% of chromatin-associated miRNAs harbor sequences fulfilling the requirements for triplex formation (Fig. 3A). More than half of these predicted triplex-forming miRNAs were shared between whole and active chromatin fractions (Fig. 3B), indicating consistent association across chromatin contexts. Among this subset, miR-7-1, miR-454, and miR-337 were identified as the most highly enriched (Figure 3C).

**Figure 3.**
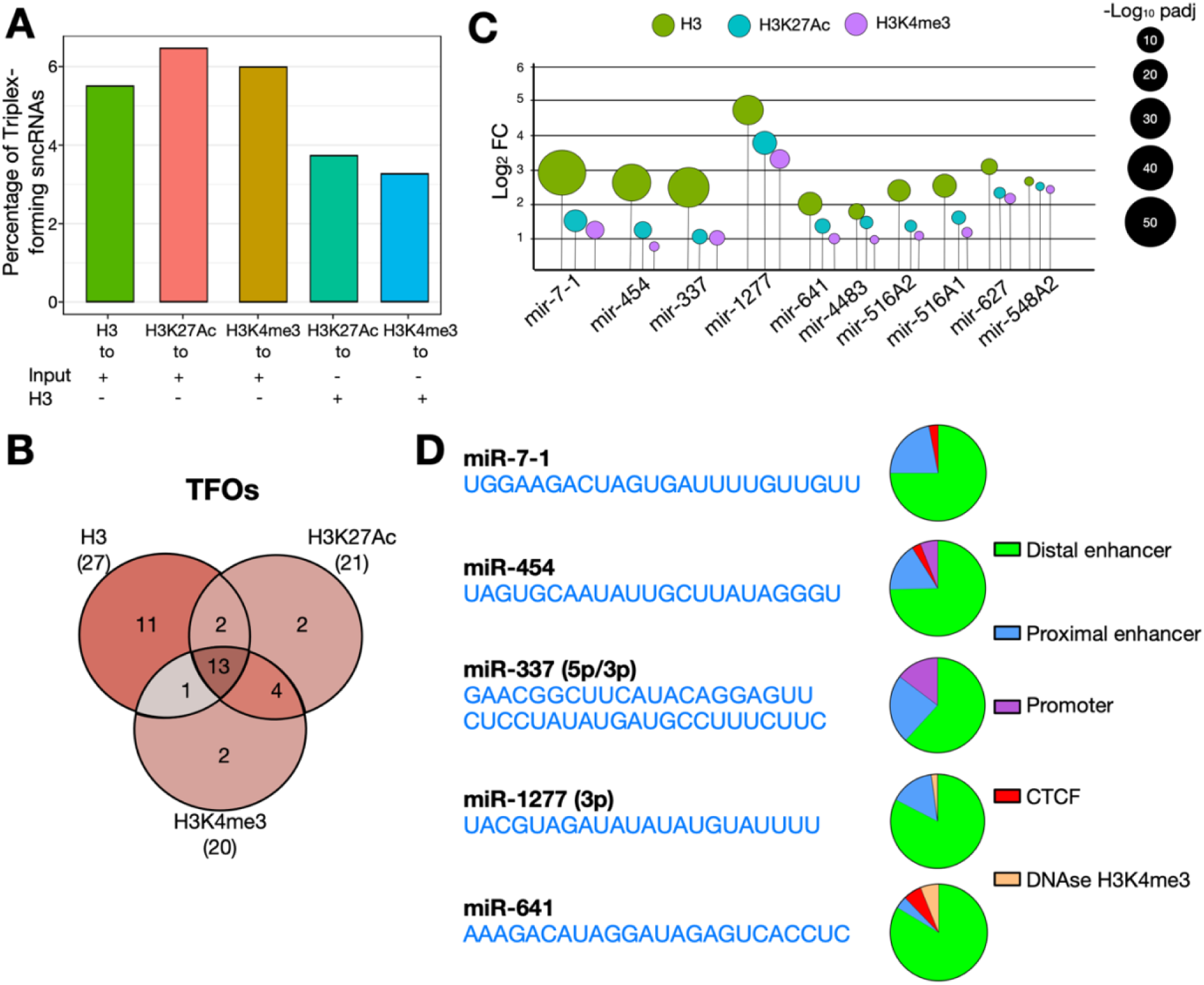
Triplex-forming miRNAs associated with chromatin in PANC-1 cells. A) Percentage of small ncRNAs harboring predicted triplex-forming sequences in each fraction, compared to nuclear or chromatin-associated ncRNAs as indicated. B) Venn diagram showing shared and specific triplex-forming small ncRNAs across samples. C) Lollipop plot displaying enrichment significance (log2 FC and -log10 adjusted *p*-value) of the most enriched triplex-forming miRNAs relative to nuclear small ncRNAs. D) Representative sequence (left) and genomic distribution of predicted triplex target sites (TTSs) (right, pie chart) for the most enriched triplex-forming miRNAs.

To provide further context, the predicted TTSs were distributed across hundreds of genomic loci, with no evidence of dominance by a small number of repetitive or polypurine-rich regions; instead, the candidate sites were enriched at regulatory elements including enhancers and promoters (Fig. 3D). These observations identify a distinct subset of chromatin-associated miRNAs with sequence features compatible with triplex-mediated chromatin targeting.

### Argonaute 2 directly binds triple helix structures

Because nuclear miRNAs commonly function in association with Argonaute proteins, and that Ago2 and other components of the RISC complex have previously been detected in the nuclear compartment (9,39), we investigated whether Ago2 can interact with ssRNA-dsDNA triplex structures.

Fluorescent electrophoretic mobility shift assays (EMSAs) were performed using pre-formed triplex substrates composed of Cy5-labeled dsDNA (TTS) and Cy3-labeled ssRNA triplex-forming oligonucleotides (TFO). In the absence of protein, a stable triplex signal was detected as a merged Cy5/Cy3 fluorescent band. Here it is important to note that the ssRNA-TFO runs at a very similar level than the TTS-TFO triplex complex, however, triplex formation is clearly showed by the TTS bandshift at lane 3 in the red channel (Fig. 4A). Increasing concentrations of recombinant Ago2 (lanes 10 to 4) produced a progressive mobility shift of the Cy5/Cy3 fluorescent triplex band, indicating formation of Ago2-triplex complexes (Fig. 4A). Notably, the shift of the triplex signal occurred at lower protein concentrations than the shift observed for the TTS, suggesting a preferential interaction of Ago2 with triple-helical structures compared to dsDNA (Figure 4A).

**Figure 4.**
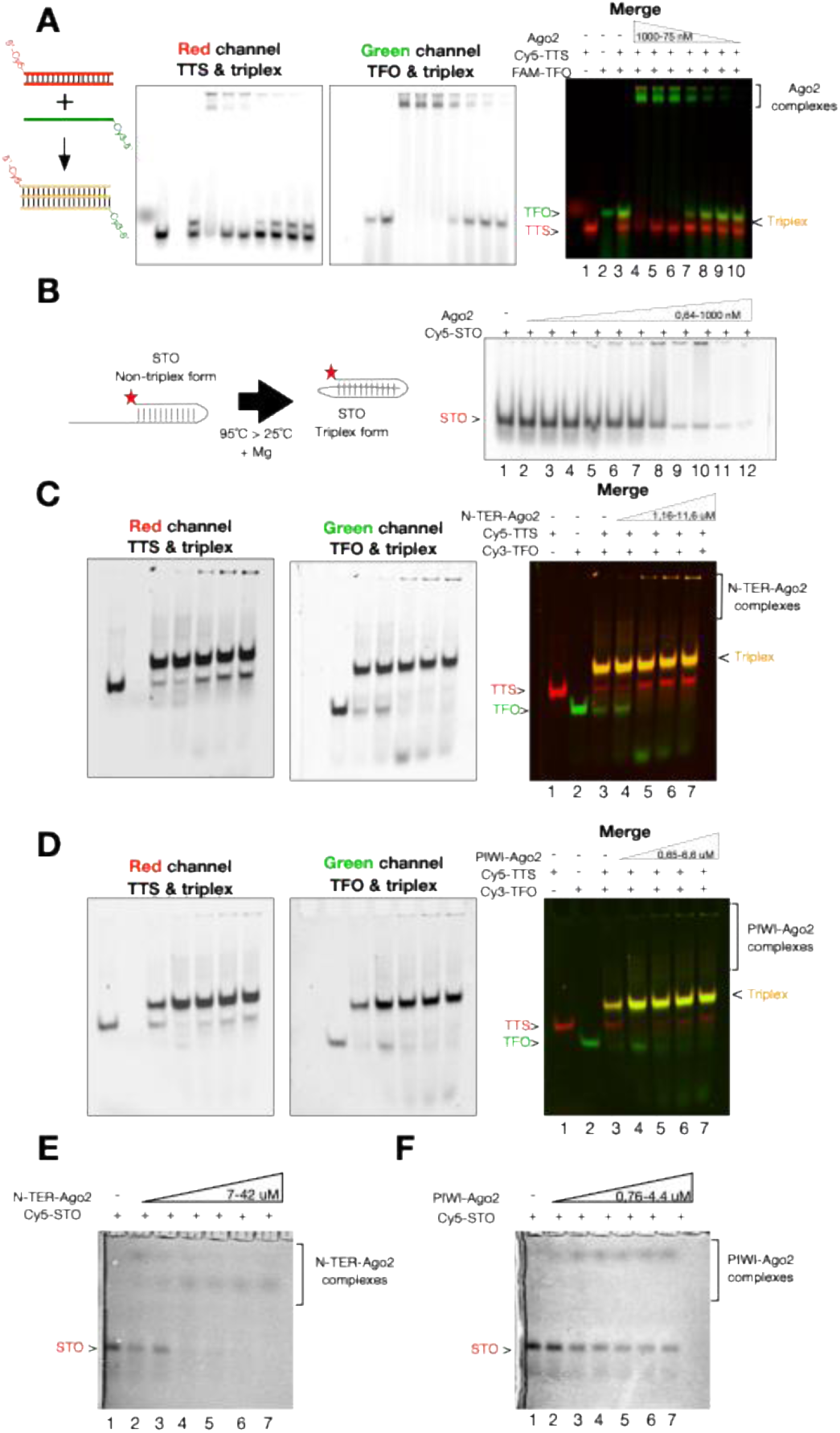
Argonaute 2 interacts with triple helices. A) A scheme depicting triplex formation with fluorophore positions is shown on the left. EMSA with triplexes formed using a fixed ratio of Cy5-labeled dsDNA-TTS and Cy3-labeled ssRNA-TFO, followed by incubation with increasing concentrations of full-length Argonaute-2 (Ago2). Lanes 1–3 correspond to dsDNA-TTS, ssRNA-TFO, and pre-assembled triplex controls, respectively. Cy5 (red), Cy3 (green), and merged channels are shown to visualize dsDNA-TTS, ssRNA-TFO, and triplex structures as indicated. B) Left, schematic representation of the self-triplex-forming oligonucleotide (STO). Right, EMSA with a fixed amount of Cy5-labeled STO incubated with increasing concentrations of full-length Ago2. C, D) EMSA with fluorescent triplexes (Cy5-labeled dsDNA-TTS and Cy3-labeled ssDNA-TFO) incubated with increasing concentrations of the Ago2 N-terminal (N-TER) domain (C) or PIWI domain (D). E, F) EMSA with Cy5-labeled STO incubated with increasing concentrations of the Ago2 N-terminal (N-TER) domain (E) or PIWI domain (F). All protein concentrations used on each assay are shown inside the corresponding triangle.

To minimize the interference of the free ssRNA-TFO with the triplexes signal observed in Figure 4A, we designed a self-triplex-forming oligonucleotide (STO) that assembles into a stable triple helix (Fig. 4B, scheme at the left). Incubation of the STO with increasing concentrations of Ago2 similarly resulted in a concentration-dependent mobility shift (Fig. 4B, gel at the right, lanes 6-12), confirming a direct interaction between Ago2 and triple-helical structures.

Ago2 comprises multiple domains that differentially mediate interactions with target mRNAs, N-Terminal and PAZ domains, and miRNA-mRNA target duplexes, including MID, PIWI, L1, and L2 domains (40,41). To explore the structural basis for the triplex interaction, we cloned, expressed, and purified recombinant GST-tagged proteins corresponding to one of each type of interaction, the N-terminal and PIWI domains of Ago2 (Supplementary Figure 3A-D).

Triplex-binding assays were performed using fluorescently labeled TTS and TFO oligonucleotides, both in DNA form to improve triplex resolution. Increasing concentrations of either domain induced a mobility shift of the triplex signal, with the N-Terminal domain producing a stronger effect than the PIWI domain (Figures 4C and D). Notably, increasing concentration of both domains also enhanced the signal corresponding to TTS alone (increase of the red band, lanes 5-7, Figures 4C and D), suggesting triplex dissolution by TFO interaction. Then, to selectively evaluate domain-specific interactions with triple helices, we repeated the assays using the STO substrate. In agreement with previous results, both the N-Terminal and PIWI domains induced concentration-dependent shifts of the STO band, again with a weaker effect observed for the PIWI domain (Figures 4E and F, lanes 2-7). The EMSA controls incubating the GST tag with triplexes and the recombinant proteins with the TTS or TFO, demonstrate that there is no interaction between the GST tag and the triplexes (Supplementary Figure 4A), and that the recombinant proteins interact with the TFO but do not with the TTS (Supplementary Figure 4B-C).

Together, these results demonstrate that Ago2 directly interacts with triple helix structures, ssRNA-dsDNA or ssDNA-dsDNA, *in vitro*. The changes in band intensities at higher protein concentrations suggest that Ago2 binding at elevated concentrations may partially destabilize the triplex by interacting with the TFO strand and promoting its dissociation from the TTS, a behavior that could be functionally relevant if Ago2 acts to resolve rather than stabilize triplexes in a cellular context. We acknowledge that these experiments do not directly demonstrate triplex formation between specific endogenous chromatin-associated miRNAs and their predicted genomic target sites (mainly due to unstable triplex formation *in vitro*); this constitutes an important limitation of the current study and a key direction for future investigation.

### Triplex-forming chromatin miRNAs show restricted evolutionary distribution

The ability of Ago2 to associate with triple-helical structures *in vitro* suggests that specific sequence features may underlie the chromatin association of a subset of miRNAs. Because triplex formation depends on specific nucleotide composition and sequence organization, these features would be expected to be under evolutionary constraint if they contribute to miRNA function. We therefore investigated the evolutionary conservation of the sequence and the phyletic distribution of chromatin-associated miRNAs (Figure 5). Our comparative analysis across jawed vertebrates revealed striking differences in the phyletic distribution of triplex-forming and non-triplex miRNAs (Figure 5). Triplex-forming miRNAs display a restricted phyletic distribution, with most members confined to therian mammals (placentals and marsupials) (Figure 5). An exception is miR-7-1, which is present in all sampled species and shows complete sequence conservation over approximately 462 million years of evolution (Figure 5). Notably, the full complement of triplex-forming miRNAs is restricted to anthropoid primates, the clade comprising apes, Old World monkeys, and New World monkeys (Fig. 5). In contrast, non-triplex-forming miRNAs display a markedly broader distribution, being present across nearly all sampled species with only a few lineage-specific absences (Figure 5).

**Figure 5.**
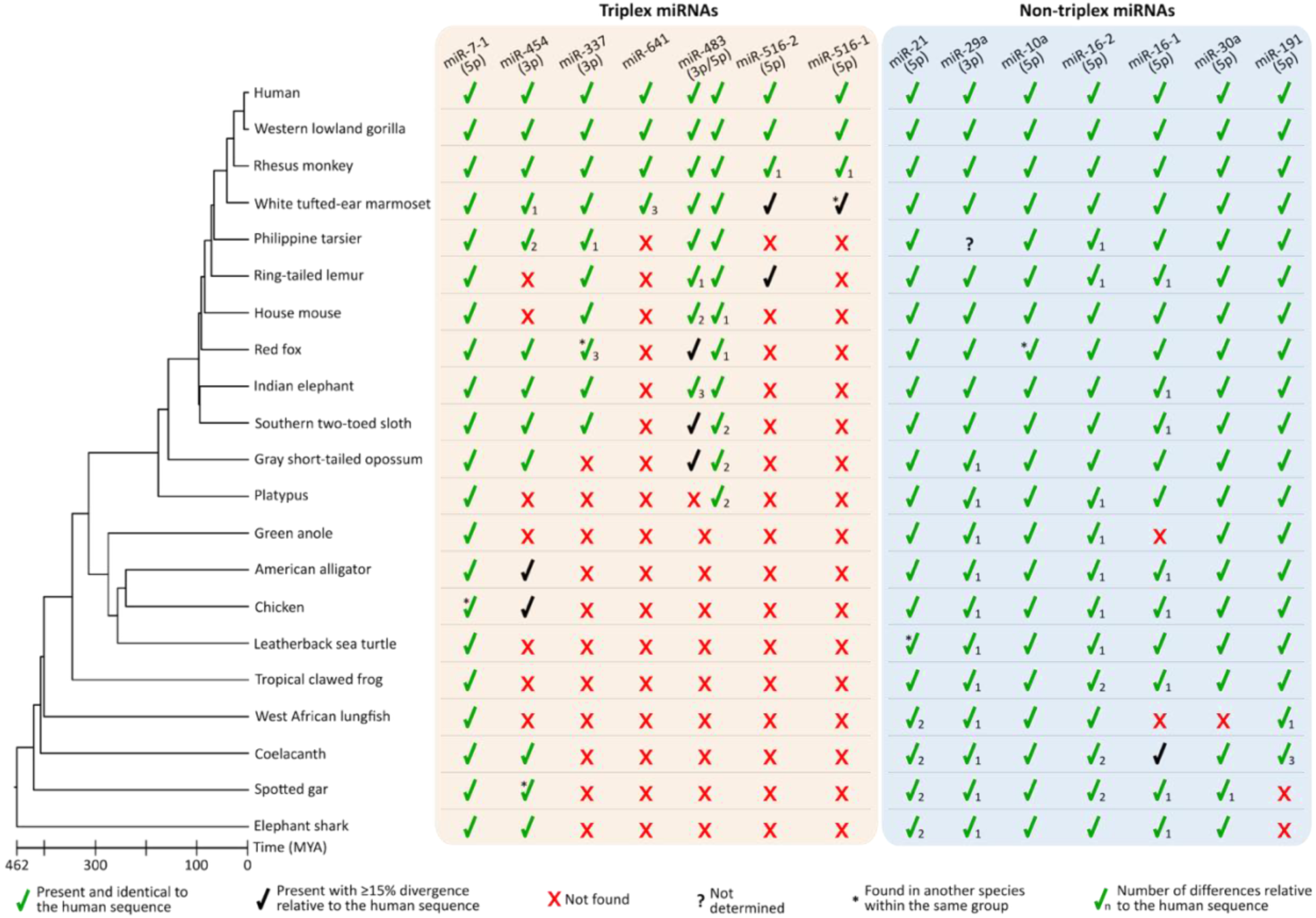
Evolutionary distribution and conservation of triplex-forming and non-triplex miRNAs across jawed vertebrates. The time-calibrated phylogeny on the left depicts the evolutionary relationships of the sampled species (MYA, million years ago). The middle panel shows miRNAs predicted to form ssRNA-dsDNA triplexes (“Triplex miRNAs”), whereas the right panel displays miRNAs lacking triplex-forming capacity (“Non-Triplex miRNAs”).

## Discussion

Our study identifies a defined population of miRNAs associated with chromatin in PANC-1 cells and provides biochemical insight into how these small RNAs may engage genomic DNA. While miRNAs are best known for their cytoplasmic roles in post-transcriptional gene regulation, increasing evidence indicates that a subset of miRNAs localizes to the nucleus and participates in chromatin-associated processes (42–44). Our results support this emerging view by demonstrating that chromatin-associated small RNAs represent a distinct population enriched in miRNAs, rather than a mere reflection of the bulk nuclear RNA pool.

To enrich specifically for transcriptionally active chromatin, we used antibodies against H3K27ac and H3K4me3, which mark active enhancers and active gene promoters, respectively, in addition to the pan-nucleosomal H3 antibody. We acknowledge that our experimental design did not include repressive chromatin marks such as H3K9me3 or H3K27me3, which would have provided an important biological contrast, allowing an assessment of whether the enrichment of miRNAs is specifically associated with active, rather than chromatin in general. This remains an important limitation and will be addressed in future work. Nevertheless, the preferential association of the predicted triplex target sites with enhancers and promoters (Fig. 3D) is consistent with a bias toward active regulatory regions, even if this cannot be conclusively established from the current immunoprecipitation data alone.

Among these chromatin-associated miRNAs, miR-21 emerged as the most abundant species. MiR-21 is one of the most extensively studied miRNAs and plays prominent roles in cancer, inflammation, and stress responses through post-transcriptional repression of numerous target transcripts (45,46). The strong chromatin abundance suggests that miR-21 may also participate in nuclear regulatory processes. However, due to its sequence, miR-21 was not identified among the subset of miRNAs with predicted triplex-forming capacity. Its chromatin association is therefore likely mediated by alternative mechanisms, most plausibly recruited as part of Argonaute-containing ribonucleoprotein complexes to chromatin-associated RNA targets, consistent with recent evidence for nuclear Ago2:miRNA activity on chromatin-associated transcripts (39). The co-occurrence of strong chromatin enrichment for miR-21 and triplex-forming potential for a distinct miRNA subset, reflects the mechanistic diversity of miRNA-chromatin interactions and should not be interpreted as a single unified mechanism. We acknowledge that this represents two partially independent observations within the study, and that experimentally linking them constitutes an important next step.

A key mechanistic insight from our study is the demonstration that Ago2 directly interacts with triple-helical nucleic acid structures *in vitro*. These experiments used previously tested triplex substrates and recombinant Ago2, thus, they do not demonstrate that specific endogenous chromatin-associated miRNAs form triplexes at their predicted genomic target sites in cells. This constitutes a central limitation of the current work, mainly due to the instability of such sequences for *in vitro* studies like EMSAs. Demonstrating the sequence-specific miRNA-genomic DNA triplex interactions in a cellular context, for example, by mutating predicted triplex-forming sequences within specific miRNAs and examining the impact on chromatin association, or by combining Ago2 CLIP with triplex-specific sequencing approaches, will be essential to validate the proposed model. Nevertheless, the biochemical evidence that Ago2 can associate with triple-helical structures, combined with the computational identification of chromatin-associated miRNAs with triplex-forming sequence features, provides a coherent framework for investigating this mechanism further.

The domain-level EMSA analyses suggest that both the N-terminal and PIWI domains of Ago2 contribute to triplex recognition, though the observed shifts for the isolated domains are modest compared to the full-length protein. This is consistent with the expectation that individual domains may contribute partially to an interaction that is strengthened by the intact multidomain architecture of Ago2. The observed concentration-dependent changes in band intensities, including apparent triplex destabilization at high protein concentrations, are interesting and could reflect an active role for Ago2 in resolving as well as recognizing triple-helical structures. However, these initial approaches lead to the exploration of quantitative affinity measurements or competition assays and further structural and biophysical studies.

Finally, our evolutionary analyses reveal a striking contrast in the phyletic distribution of triplex-forming versus non-triplex-forming chromatin-associated miRNAs. Non-triplex-forming chromatin miRNAs are broadly conserved across jawed vertebrates, consistent with their roles in conserved regulatory processes. In contrast, the full complement of triplex-forming chromatin-associated miRNAs is restricted to anthropoid primates, except for miR-7-1, which is deeply conserved and shows complete sequence conservation over approximately 462 million years of evolution. This evolutionary pattern suggests that triplex-mediated miRNA-chromatin interactions represent a relatively recent regulatory innovation that emerged during primate evolution, potentially contributing to the expanded transcriptional complexity associated with this lineage (47,48). This finding is consistent with broader observations that lineage-specific miRNAs frequently contribute to the fine-tuning of gene regulation in a species-specific manner (49–52).

Together, our findings contribute to the growing body of evidence supporting chromatin-associated functions for miRNAs, a regulatory dimension that remains insufficiently explored. Our data reveal that chromatin-associated small RNAs in PANC-1 cells are enriched in miRNAs, where miR-21 is the most abundant, and that a subset of these harbor sequence features compatible with RNA-DNA triplex formation. In parallel, we show biochemically that Ago2 can interact with triple-helical nucleic acid structures *in vitro*. These observations are suggestive rather than mechanistically conclusive, they do not yet establish that endogenous chromatin-associated miRNAs form sequence-specific triplexes at genomic loci, nor that Ago2 is recruited to chromatin through this mechanism in cells. We present them as a coherent set of descriptive and biochemical data that together motivate a testable hypothesis and hope they will stimulate further experimental work on the nuclear and chromatin-level functions of small non-coding RNAs.

## Declaration of interest

The authors declare that they have no conflict of interest

## Data availability

Small RNA sequencing from the chromatin-RNA immunoprecipitation assays data has been deposited in NCBI’s Gene Expression Omnibus (GEO) and is accessible through GEO series accession number GSE320385.

## Acknowledgments

This work was supported by the grants from Agencia Nacional de Investigación (ANID) FONDEF 25I10174 (RMal), FONDECYT Iniciación 11230662 (RMun), FONDECYT Regular 1251655 (GM), 1250688 (JCO), FONDECYT Exploración 13250135 and Bio & Medical Technology Development Program of the National Research Foundation (NRF) funded by the Korean government (MSIT) (No. RS-2025-15373195) (GVN).

## Declaration of generative AI and AI-assisted technologies in the manuscript preparation process

During the preparation of this work, the author(s) use Perplexity for language edition. The author(s) reviewed and edited the output as needed and take full responsibility for the content of the published article.

## Notes

### Competing Interest Statement

The authors have declared no competing interest.

### Summary of Updates

The Abstract, Introduction, Results, and Discussion has been updated for clrarifucation; Figure 2 revised, and supplemental files has been added.

