## Supplementary Figure 1 for "RNA-DNA triplex-forming miRNAs define an evolutionarily recent chromatin regulatory mechanism"

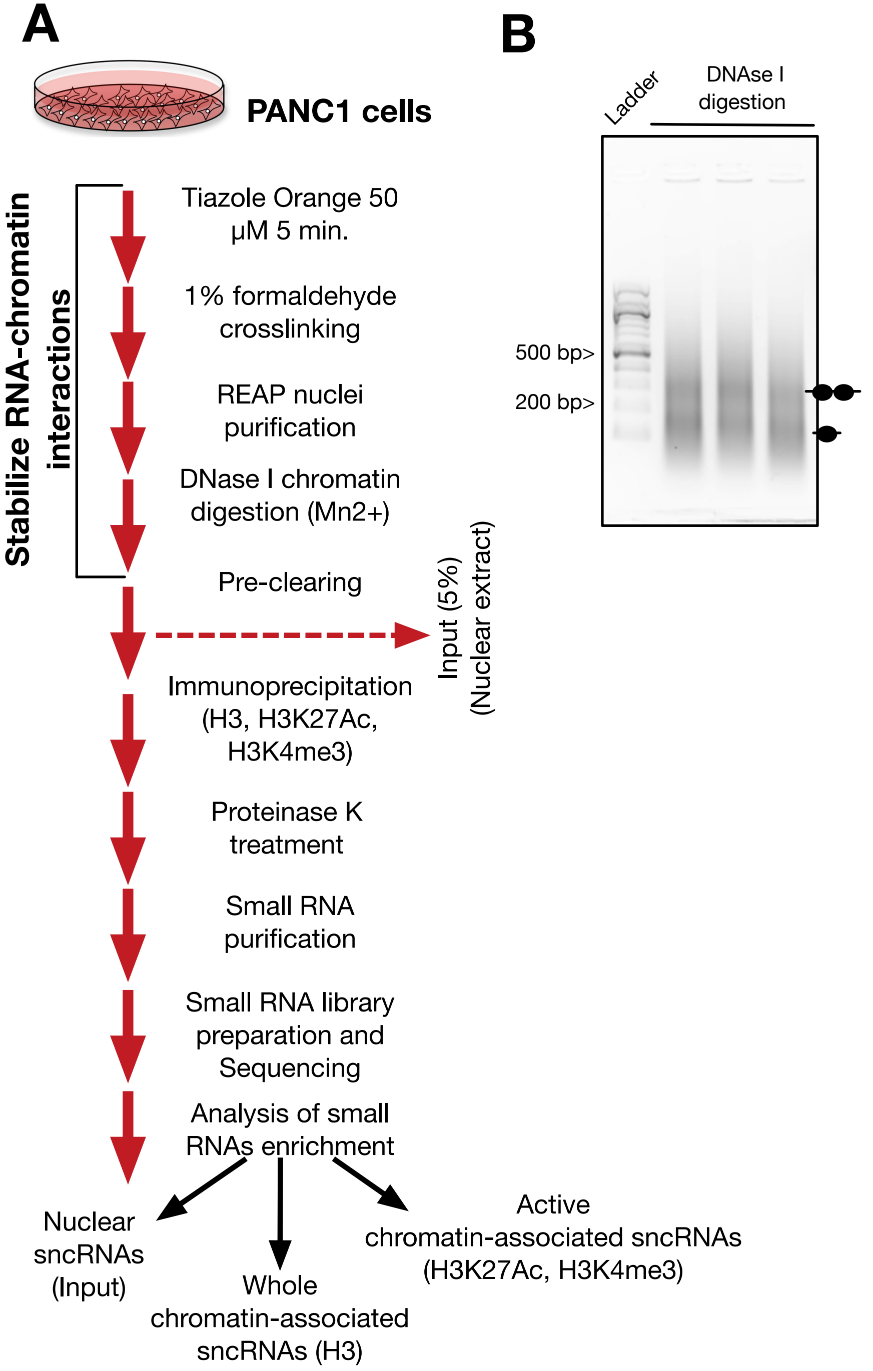


**Supplementary Figure 1. Chromatin-RNA immunoprecipitation for small RNA sequencing.** A) Step-by-step diagram showing all the procedures performed to sequence chromatin-associated and bulk nuclear small RNAs of PANC-1 cells. B) DNA size distribution after DNAse I chromatin digestion under the conditions indicated on A).
