## Supplementary Figure 2 for "RNA-DNA triplex-forming miRNAs define an evolutionarily recent chromatin regulatory mechanism"

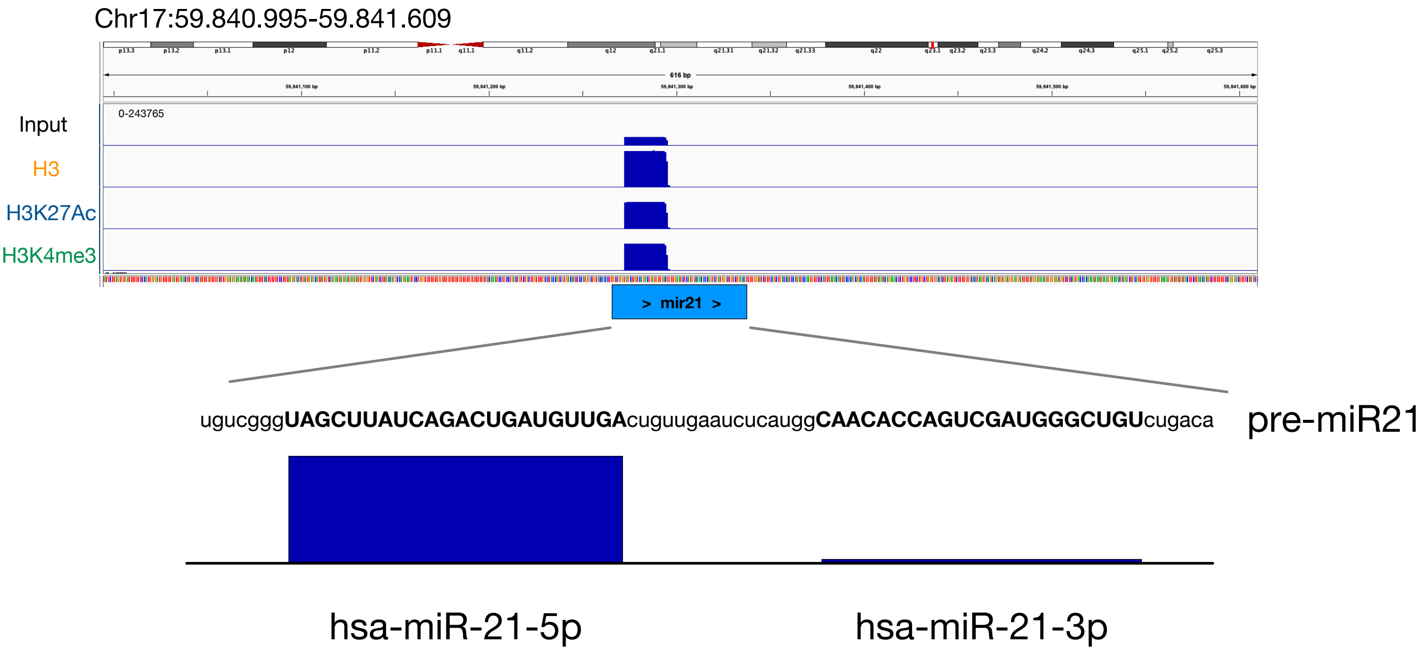


**Supplementary Figure 2. The mature form of miR21 hsa-miR-21-5p, and not the -3p, is the most abundant in chromatin.** IGV scheme depicting the sequencing reads over the human miR21 gene in all the samples analyzed.
