## Supplementary Figure 3 for "RNA-DNA triplex-forming miRNAs define an evolutionarily recent chromatin regulatory mechanism"

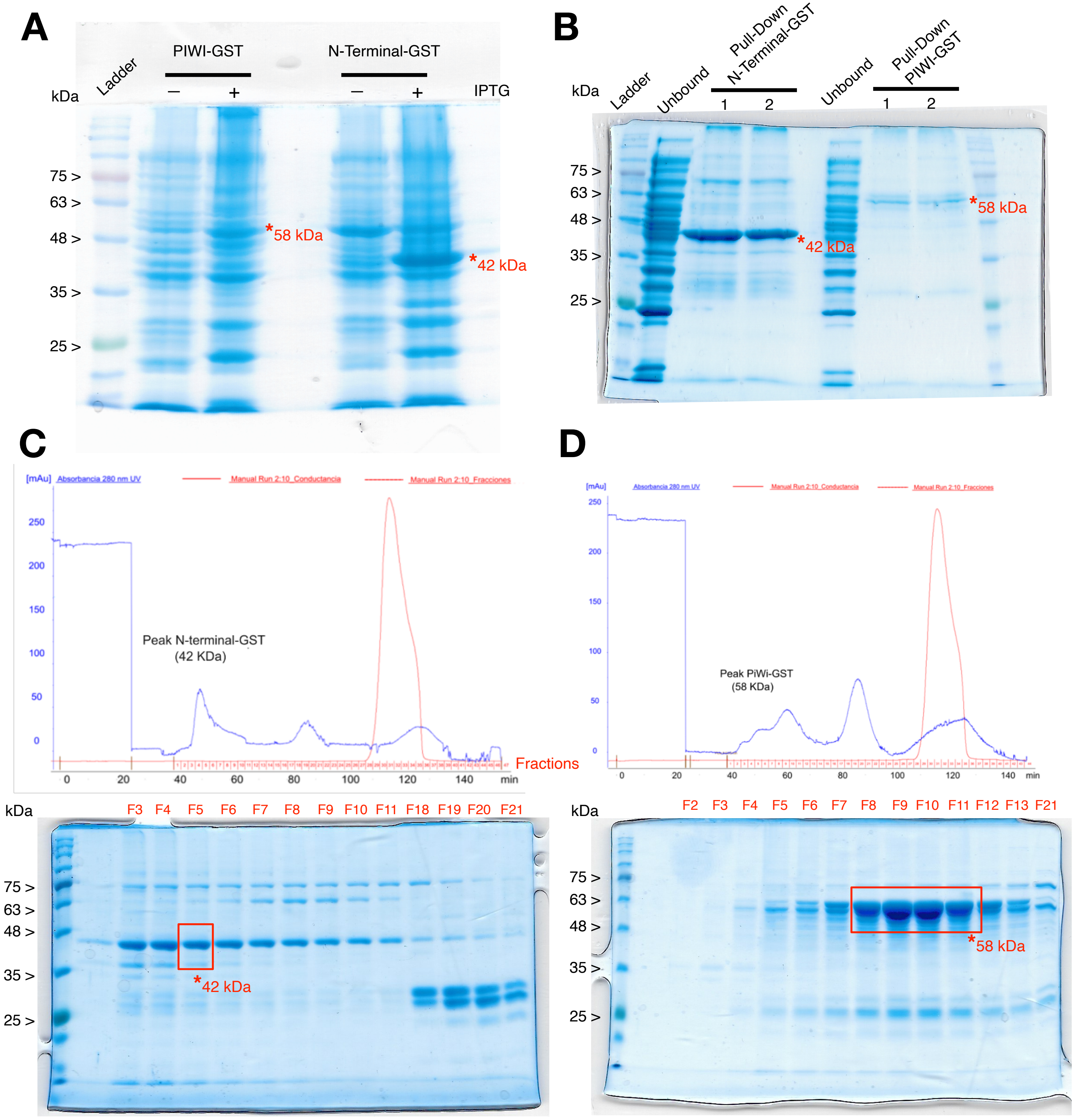


**Supplementary Figure 3. Expression and purfication of Ago2 N-terminal- and PIWI-GST recombinant proteins.** A) IPTG induction of recombinant proteins in *Escherichia coli* B834(DE3) p-LyS cells, analyzed by SDS-PAGE at 4 hours post-induction. The asteriscs show the predicted sizes for each recombinant protein. B) After induction, glutathione affinity purification of the recombinant proteins from large scale cultures. Unbound correspond to the fractions of the samples that were not bound to the gluthatione sepharose beads. C) and D) Size exclusion chromatography to purify the N-Terminal-GST (C) and PIWI-GST (D) recombinant proteins. The chromatogram monitoring proteins at 280 nm is shown in the upper pannels (blue lines), while fractions intervals are shown above the X axis in red. Fractions including the main peaks were analyzed by SDS-PAGE. The read boxes indicate the samples that were pooled and concentrated for the EMSAs.
