## Supplementary Figure 4 for "RNA-DNA triplex-forming miRNAs define an evolutionarily recent chromatin regulatory mechanism"

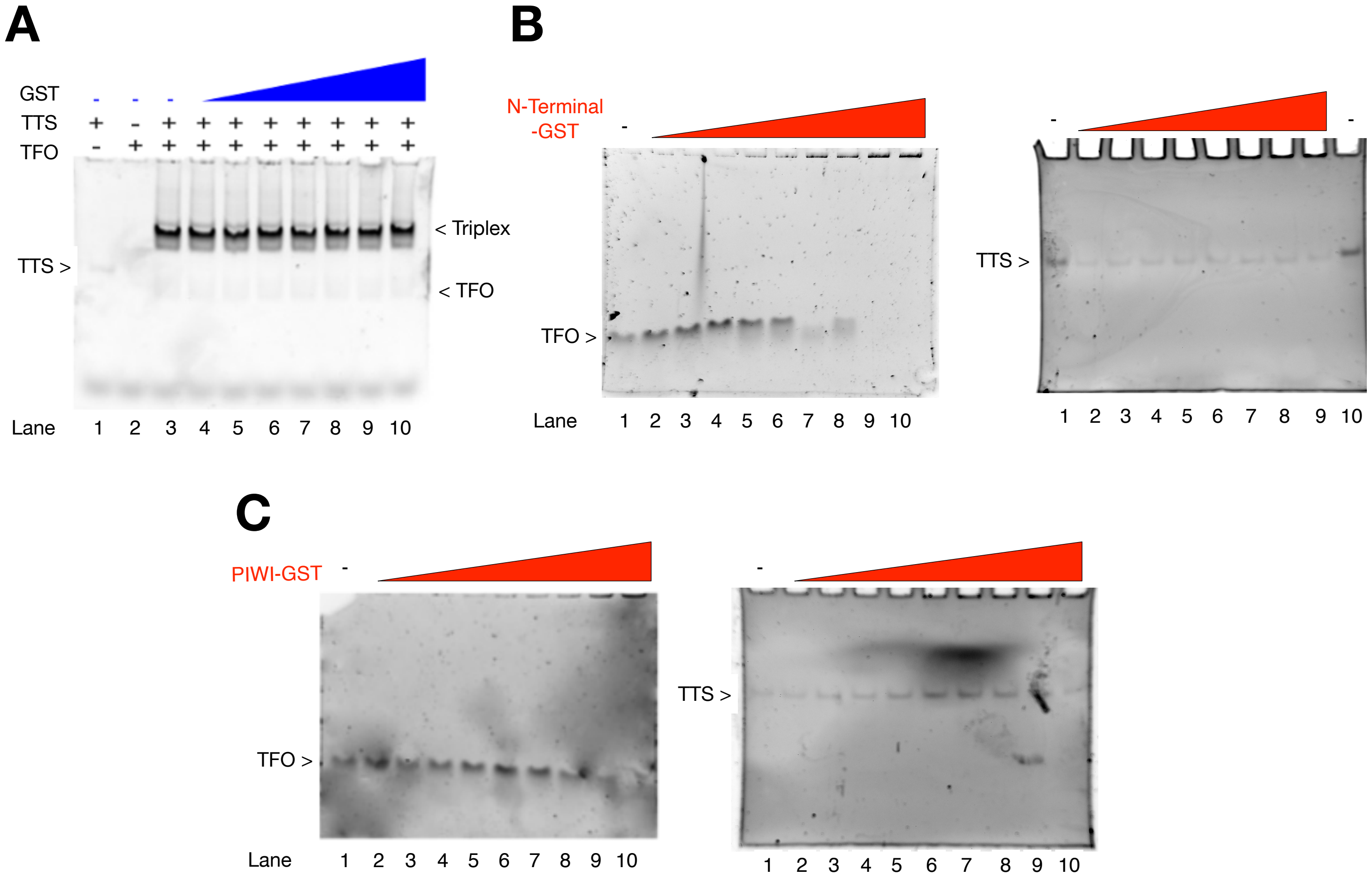


**Supplementary Figure 4. Recombinant protein interaction with triplexes and their constituents. A)** Non-labeled TTS (lane 1) and TFO (lane 2) were mixed on a 1:2 ratio to form a non-labeled triples (lane 3). Then, EMSA was performed by incubating increasing concentration of GST tag (50-900 nM) with non-labeled triplexes (lanes 4 to 10). **B) and C)** EMSAs where non-labeled TFO (lane 1, gel at left) and non-labeled TTS (lane 1, gel at right) were incubated with increasing concetrations of Ago2 N-terminal-GST (lanes 2-10) (B) or PIWI-GST (C). All these assays were electrophresed on 12% polyacrlamide gels prepared on TA buffer. Non-labeled triplex, TTS and TFO were visualized after incubation with 50 nM Tiazole orange for 10 minutes on TA buffer.
